# Interior pH Sensing Residue of Human Voltage-Gated Proton Channel H_v_1 is Histidine 168

**DOI:** 10.1101/2022.12.07.519452

**Authors:** Mingzhe Shen, Yandong Huang, Zhitao Cai, Vladimir V. Cherny, Thomas E. DeCoursey, Jana Shen

**Author notes:** Joint first author.

## Abstract

The molecular mechanisms governing the human voltage-gated proton channel hH_v_1 remain elusive. Here we used membrane-enabled hybrid-solvent continuous constant pH molecular dynamics (CpHMD) simulations with pH replica exchange to further evaluate the recently obtained structural models of hH_v_1 in hyperpolarized (closed channel) and depolarized (open channel) states (Geragotelis, Tobias et al., Proc. Natl. Acad. Sci. USA 2020) and explore potential pH-sensing residues. The CpHMD titration at a set of symmetric pH conditions revealed three residues that can gain or lose protons upon channel depolarization. Among them residue H168 at the intracellular end of the S3 helix switches from the deprotonated to the protonated state and its protonation is correlated with the increased tilting of the S3 helix during the transition from the hyper-to the depolarized state. Thus, the simulation data suggest H168 as an interior pH sensor, in support of a recent finding based on electrophysiological experiments of H_v_1 mutants (Cherny, DeCoursey et al., J. Gen. Physiol. 2018). Our work represents an important step towards deciphering the pH-dependent gating mechanism of hH_v_1.

**SIGNIFICANCE:** The human voltage-gated proton channel hH_v_1 is comprised of a proton-selective voltage sensing domain and responsible for cellular pH homeostasis. Despite intense experimental and theoretical investigations, its pH-dependent gating mechanism is not understood. Our simulation data offer strong evidence supporting the role of H168 as a pH_*i*_ sensor (Cherny, DeCoursey et al., J. Gen. Physiol. 2018). Deciphering the interior pH sensor moves us a step closer to elucidating the structure-function relationship of hH_v_1.

## INTRODUCTION

The voltage-gated proton channel H_v_1 belongs to a super family of proteins which become activated through depolarization via a voltage-sensing domain (VSD) (1–3). However, unlike voltage-gated sodium or potassium channels, H_v_1 does not possess a separate ion-conduction pore domain and it is also gated by pH (4, 5). The primary function of H_v_1 is the extrusion of protons from cells to maintain pH homeostasis, but it is also involved in other biological events such as the production of reactive oxygen species, sperm mobility, and B-cell activation (see recent reviews (6, 7)). Three independent studies showed that H_v_1 is a homodimer comprising four transmembrane helical segments (S1–S4), but each monomer can function independently through the VSD which also serves as a pore (8–10). Since the discovery of the mammalian H_v_1 gene in 2006 (2, 3), the mechanisms of gating and proton selectivity of human H_v_1 (hH_v_1) have been extensively studied using electrophysiology and mutagensis (5, 11).

H_v_1 is closed under a hyperpolarizing membrane voltage, i.e., the voltage is more negative inside the cell as compared to outside, and it opens upon membrane depolarization. A unique property of H_v_1 is that its opening voltage is dependent on the pH gradient across the membrane, which is a phenomenon known as ΔpH dependent gating (6). The proton conductance-voltage (*g*_H_–*V*) relationship shifts negatively by 40 mV with a one-pH unit increase of the outside pH (pH_*o*_) or a one-pH unit decrease of the inside pH (pH_i_) (6), which is commonly known as the rule of forty. A recent electrophysiological study (12) demonstrated that the mutation of H168 to Q168, T168, or S168 compromises the pH_i_ dependence of the gating of hH_v_1.

Despite enormous progress, the detailed molecular mechanism of H_v_1 gating remains elusive due to limited structural information. At present, two atomic-resolution experimental structure models are available for the resting or closed state and for an intermediate state of H_v_1 (13, 14), but an experimental structure model for the depolarized open state is lacking. The model representing the closed state of H_v_1 is an X-ray crystal structure of a chimeric construct of the mouse H_v_1 (mH_v_1cc) at 3.45 Å resolution, which shows a monomeric antiparallel four-helix-bundle (S1 through S4) arranged in in the shape of “a closed umbrella” (PDB: 3WKV) (13). The voltage-sensing S4 contains three gating arginines, R201, R204, and R207, corresponding to R205, R208, and R211 in hH_v_1, respectively. We will refer to them as R1, R2, and R3, respectively hereafter. While R1 and R2 form salt bridges with D108 (D112 in hH_v_1), R3 forms a salt bridge with D170 (D174 in hH_v_1). D112 (in hH_v_1) is crucial to proton selectivity (5). H136 (H140 in hH_v_1) interacts with E115 and D119, which are respectively E119 and D123 in hH_v_1. Residues S94, I122 to A131, P159 to F161, and K187 to H190 (residue numbering in mH_v_1cc) are not resolved, and 16 other residues have missing atoms. The model representing an intermediate state of H_v_1 comes from a solution NMR model of the VSD of hH_v_1cc embedded in mixed detergent micelles (14). Finally, an electron paramagnetic resonance (EPR) study of H_v_1 combined with the crystal structure of the *Ciona intestinalis* voltage sensing phosphatase VSD (CiVSD) (15) resulted in closed and open state models of hH_v_1 (16). Given the limited experimental structural information on H_v_1, molecular dynamics (MD) simulations starting from various homology models (17–23) have been produced to explore the dynamical properties of the channel. Recently, MD simulations (22, 23) have been performed to explore the open state of H_v_1. Starting from a homology model of monomer hH_v_1 based on the X-ray structure of mmH_v_1cc (13), Tobias and coworkers conducted MD simulations first without and then with an external field (23). In the latter portion, a constant uniform electric field corresponding to a transmembrane potential of -150 mV was applied for 5 μs followed by a switch to +150 mV for about 22 μs (23). From these simulations, Tobias and coworkers obtained the hyperpolarized (Hypol, closed) and depolarized (Depol, open) structural models of H_v_1 (Figure 1a).

**Figure 1.**
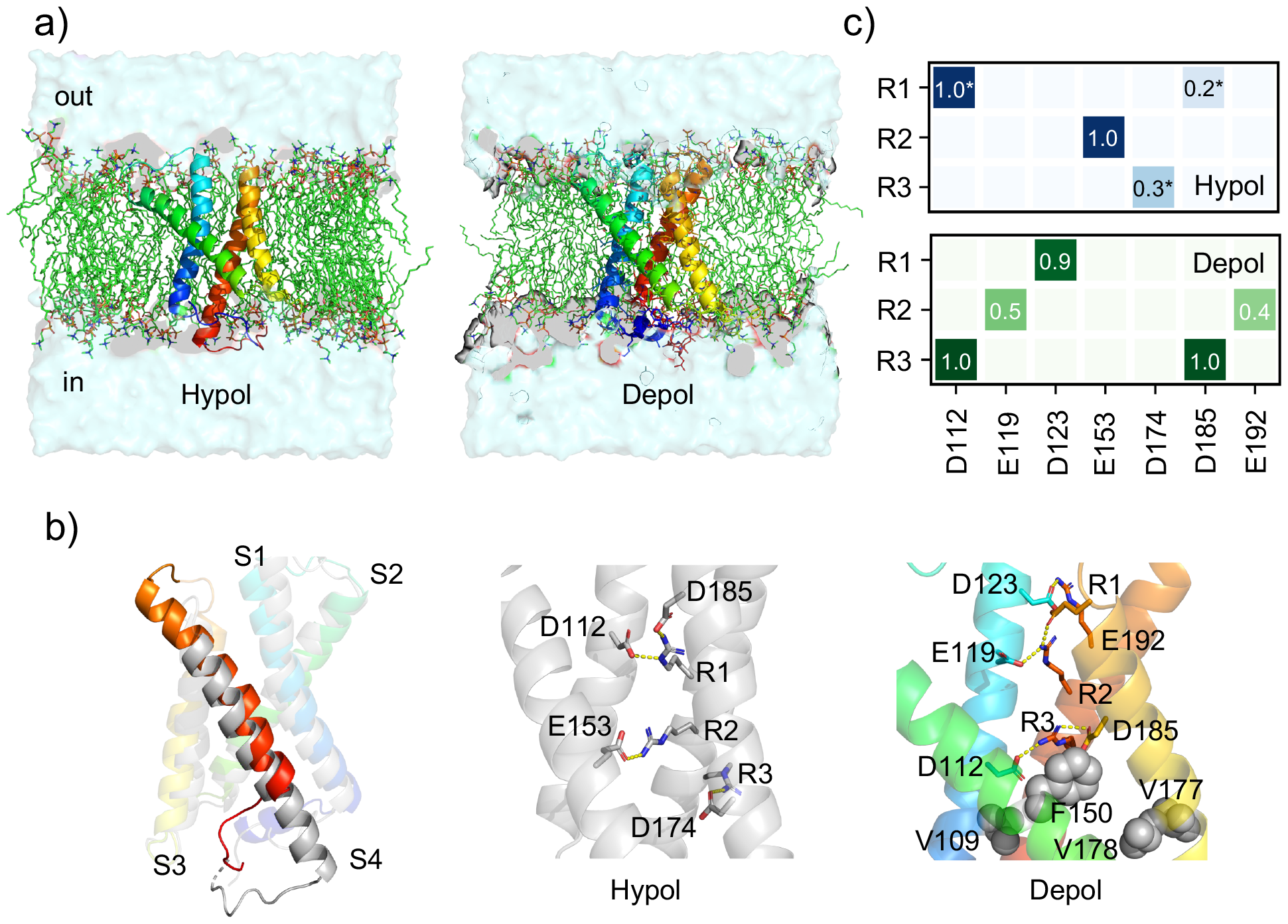
The hyperpolarized and depolarized states of hH_v_1 show different salt-bridge networks and p*K*_a_ values for three residues. **(a)** Starting structures of the hyperpolarized (Hypol, left) and depolarized (Depol, right) states of H_v_1 embedded in the solvated POPC membrane. The structures were adopted from Ref. (23). **(b)** Left. Overlay of the Hypol (grey) and Depol (rainbow colored) models (23). Middle and right are the zoomed-in views of the Hypol (middle) and Depol (right) models. Residues in the salt-bridges involving the gating arginines are shown in the stick model. The putative hydrophobic gasket residues (V109, F150, V177, and V178) (24)) are rendered as spheres. **(c)** Comparison of the salt-bridge occupancies (probabilities) in the Hypol (top) and Depol (bottom) states from the simulations of hH_v_1 at pH 7.5. A salt bridge is considered formed when the minimum distance between the carboxylate oxygen and the arginine guanidinium nitrogen is below 3.5 Å. Star indicates the agreement with the CiVSD-based resting hH_v_1 model of Li and Perozo (16).

**Figure 2.**
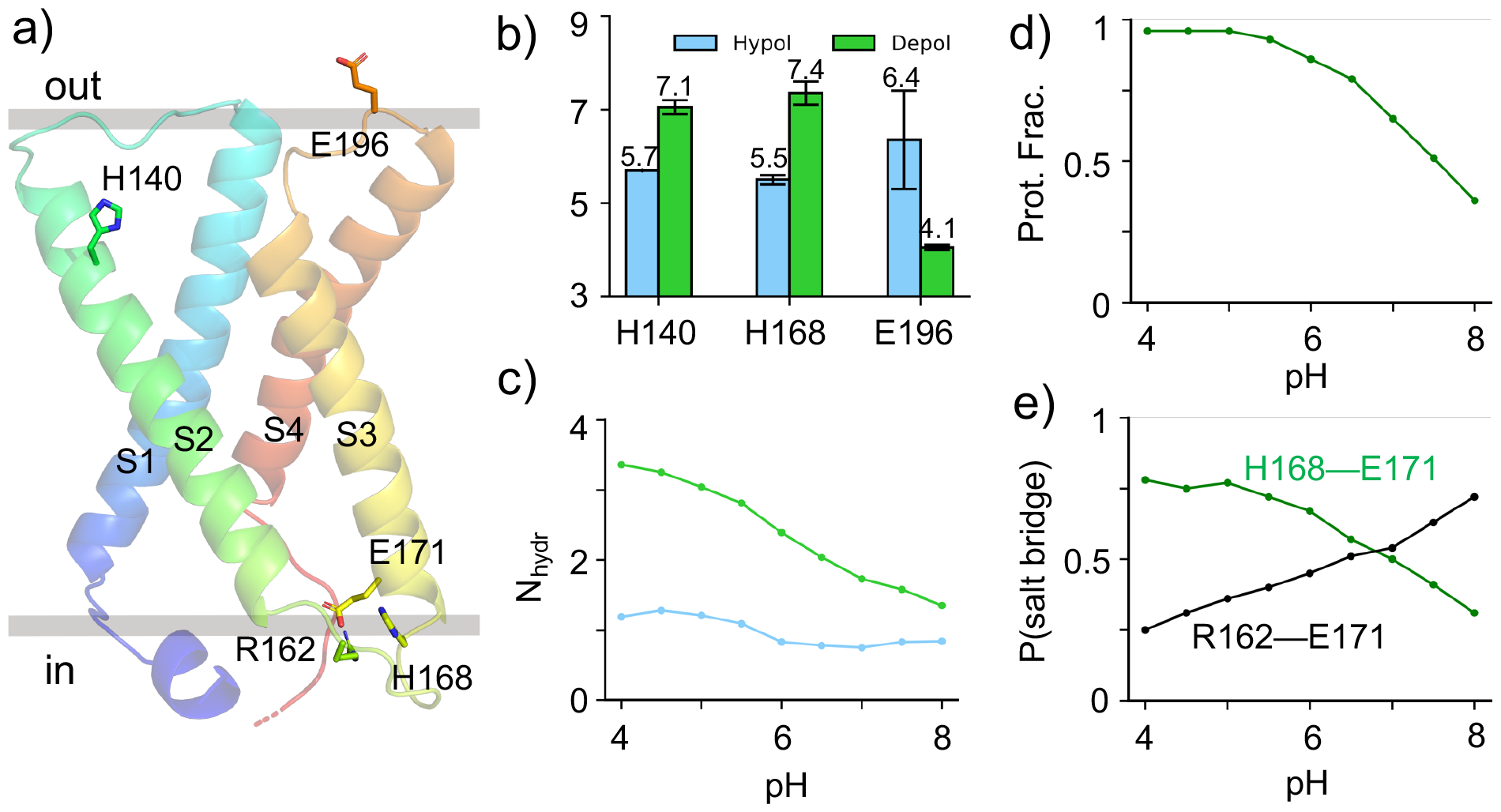
Why does H168 switch protonation state? **a)** A snapshot of hH_v_1 from the Depol state simulation. Residues discussed in the text are labeled. **b)** Calculated p*K*_a_ values of H140, H168, and E196 in the Hypol (blue) and Depol (green) states of hH_v_1. The average p*K*_a_’s from the two sets of simulations are given, and the error bars indicate the two p*K*_a_ values (Supplemental Table 1). These are the only residues that show a larger than 0.5-unit p*K*_a_ difference between the two states and may change protonation state. **c)** Number of waters (hydration number) near H168, calculated as the number of water molecules within 3.4 Å (between water oxygen and any side chain heavy atom of H168). **d)** Protonation fraction of H168 in the Depol state at different simulation pH. **e)** Probabilities (or occupancies) of the H168–E171 (green) and R162–E171 (black) salt bridges in the Depol state at different simulation pH.

In the Depol model, S4 is moved up toward the extracellular side by about 8 Å, corresponding to two helical turns or two “clicks” relative to the Hypol model (Figure 1b). This is in contrast to the S4 movement in the activation of the aforementioned Ci-VSD, which is about 5 Å or one click according to the X-ray crystal structures (15). Another dramatic conformational change in the Depol model is the S2 helix, which tilts to the side, creating a wider intracellular gap between S1 and S2. Likewise, S4 tilts away from S3, widening the intracellular gap between these two helices. Finally, S1 is moved down towards the intracellular side in the Depol model.

The objective of the present work is to further evaluate the aforementioned closed and open models from the Tobias group (23) and investigate the identities of potential pH_*i*_- and pH_*o*_-sensing residues. To accomplish this, we applied continuous constant pH molecular dynamics (CpHMD) simulations, which allows protonation states to be determined on the fly during the course of MD simulation at a specified pH condition (25). In the membrane-enabled (26) hybrid-solvent CpHMD (25), conformational dynamics of the protein is sampled in an all-atom fashion with explicit membrane and water, while the protonation states are propagated using the solvation forces calculated by a membrane generalized Born model (27). This method coupled with the pH replica-exchange protocol (25) has been validated for the determination of protonation states and proton-coupled conformational dynamics of several membrane transporters (26, 28, 29) and the M2 channel (30). Here we applied the pH replica-exchange membrane-enabled hybrid-solvent CpHMD simulations on the aforementioned Hypol and Depol models of hH_v_1 (23). Our data suggest that H168 is a pH_i_ sensor, consistent with the mutagensis and electrophysiology data of DeCoursey and coworkers (12). Our work offers an important contribution towards deciphering the pH-sensing residues in hH_v_1.

## RESULTS AND DISCUSSION

The pH replica-exchange membrane-enabled hybrid-solvent CpHMD simulations were conducted for the Hypol and Depol states of hH_v_1 starting from the respective models obtained by Tobias and coworkers based on conventional fixed-protonation-state MD simulations (Figure 1a). For each state, two independent sets of replica-exchanges CpHMD simulations were performed. The simulation set 1 was performed with 16 replicas in the pH range 3–11.5 and an aggregate sampling time of 400 ns; simulation set 2 was conducted with 24 replicas in the pH range 3–8 and an aggregate sampling time of 720 ns (see Methods). The calculated p*K*_a_’s are listed in Supplemental Table 1 and the titration data are showin in Supplemental Figure 1. Unless otherwise noted, the analysis is based on the simulation set 1 with the first 10 ns per replica of data discarded (240 ns sampling). The data of simulation set 2 are consistent with set 1 and the analysis is given in the Supplemental Figure 2–5.

### Interhelical salt-bridges involving gating arginines

The salt bridges formed between Asp or Glu and the gating Arg’s are critical for stabilizing the channel in closed or open states. It is widely believed that the interhelical salt bridges in H_v_1 are rearranged as a result of the upward movement of S4 helix when the channel switches to the depolarized state; however, the exact rearrangement or the extent of the S4 helix upshift is still under debate (6). During the simulations at different pH, most of the salt bridges in the initial Hypol and Depol models (23) (Figure 1b) are maintained with varying degree of stabilities (or occupancies) at different pH. Figure 1c shows the salt-bridge occupancies at pH 7.5. The Hypol state contains four salt bridges: those between R1 and D112 (S1) and between R2 and E153 (S2) are present in the entire duration of the simulations, while those between R1 and D185 (S3) and between R3 and D174 have occupancies of 20% and 30%, respectively (Figure 1c). Note, the R3–D174 salt bridge is absent in the initial Hypol model (the minimum distance between the Arg nitrogen and Asp oxygen is 4.7 Å) but forms during the simulations.

Comparison of the Hypol-state simulations with the CiVSD based resting model of hH_v_1 dimer of Li and Perozo (16) shows that the R1 salt-bridge interactions with D112 and D185 and the R3 salt-bridge interaction with D174 are consistent; however, the R2–E153 interaction contradicts the CiVSD based model in which E153 interacts with R3 (16). The discrepancy is due to the relative position of S2 (which harbors F150 and E153) and S4. In the closed-state model of Tobias and coworkers which is derived from mmH_v_1cc, S2 is positioned much higher such that F150 is nearly one helical turn higher than R2 and E153 interacts with R2 (23). By contrast, in the CiVSD-based closed model (16), S2 is positioned lower such that F150 is approximately on the same level as R2 and E153 interacts with R3. Another difference between the mmH_v_1cc- and CiVSD-based closed models (16) is that S3 (which harbors D174 and D185) is positioned lower with respect to S4 in the mmH_v_1cc-based model. This discrepancy may explain why R1–D185 and R3–D174 salt bridges are weak (occupancies below 50%) in the Hypol-state simulations. Due to the higher S2 and lower S3 position in the mmH_v_1cc-based closed model, F150 is above, rather than horizontally aligned with the other hydrophobic gasket (HG) residue V178.

In the Depol state simulations, due to the upward movement of S4, all the salt bridges in the Hypol state are disrupted, and five new salt bridges are formed. R1 interacts with D123 (S1) with a salt-bridge occupancy of 90%; R2 interacts with E119 (S1) and E192 (loop connecting S3 and S4) with occupancies of 50% and 40%, respectively; and R3 maintains the salt bridge with D112 (S1) and D185 (S3) in the entire simulation time (Figure 1b and c). It is worthwhile noting that, D112 is locked in a very stable salt bridge with R3, whereas it interacts with R1 in the Hypol state. Thus, the Depol model of Tobias and coworkers represents two clicks outward movement of S4.

Although the gating arginine positions in the model of Tobias and coworkers satisfy the evidence based on the experimental data on the voltage-gated opening of Na^+^ or K^+^ channels (31), the S4 movement contradicts the one-click hypothesis for hH_v_1 based on the interpretation of experimental data (18, 32) including Cys scanning and MTS modification experiments (33, 34). MD simulations based on a homology model using the Na^+^ and K^+^ channels as templates, suggest that only R1 and R2 pass through the hydrophobic gasket, while R3 is remains on the intracellular side (18). The pH dependence of gating currents is also consistent with one-click movement of S4 (35). Finally, the EPR study of Li and Perozo indicated that channel opening could be achieved by one-click movement of S4 (16).

One possible reason for the two vs. one-click discrepancy is the large external electric field applied in the simulations. The Hypol state model was obtained with an external voltage of -150 mV (23), which is larger in magnitude than the resting potential in most cells (−40 to -90 mV, depending on the cell type) (36–38). Furthermore, the voltage (150 mV) applied to obtain the Depol model (23), is much higher than is ever experienced by most cells during their lifetime. Excitable cells such as nerve and muscle cells reach roughly +50 mV at the peak of an action potential. Phagocytes reach +50 mV during the respiratory burst (39); (40). Those who use voltage-clamp methods to study H_v_1 quickly learn that the combination of slow activation and the need to use large depolarizations at some pH often results in abrupt termination of the experiment. To extend the voltage range of data collection, one may use shorter pulses for large depolarizations. Red cell membranes rupture irreversibly within about 2 s if +200 mV is applied (41).

If H_v_1 can occupy more than two states (distinct configurations adopted during gating) then -150 mV and 150 mV would likely reflect the most extreme positions. There is abundant experimental evidence that multiple distinct states exist (42–44). In living cells, the channel may typically occupy the most superficial closed state and the nearest open state. Given these considerations, it may be futile to discuss “the” closed or open state. It is more meaningful to determine which configurations of the protein conduct and which do not. A variety of experimental evidence supports the idea that movement of the S4 helix outward by a single click suffices to open the conductance pathway (16, 18, 32, 35, 45). It is entirely possible that additional depolarization can move S4 further into another open state.

Another concern with the models on which we based our calculations is that full opening of the channel was achieved within 10 μs of application of +150 mV (23). The time constant of activation of H_v_1 in human cells is ∼220 ms at +160 mV (Fig. 3 in (46)). In mammalian cells, H_v_1 opening is sigmoidal, and can be fit approximately by a delay followed by a single exponential. However, when the channel is expressed as a monomer, gating is exponential (8, 47) and the activation time constant is up to 6.6 times faster (10, 47, 48). Correcting for the fact that the simulation was conducted on a monomer still means that 10 μs after depolarization to 150 mV only 1 in 1000 channels would be open. The nature and biological significance of the changes that occurred during the limited simulations are thus difficult to interpret.

**Figure 3.**
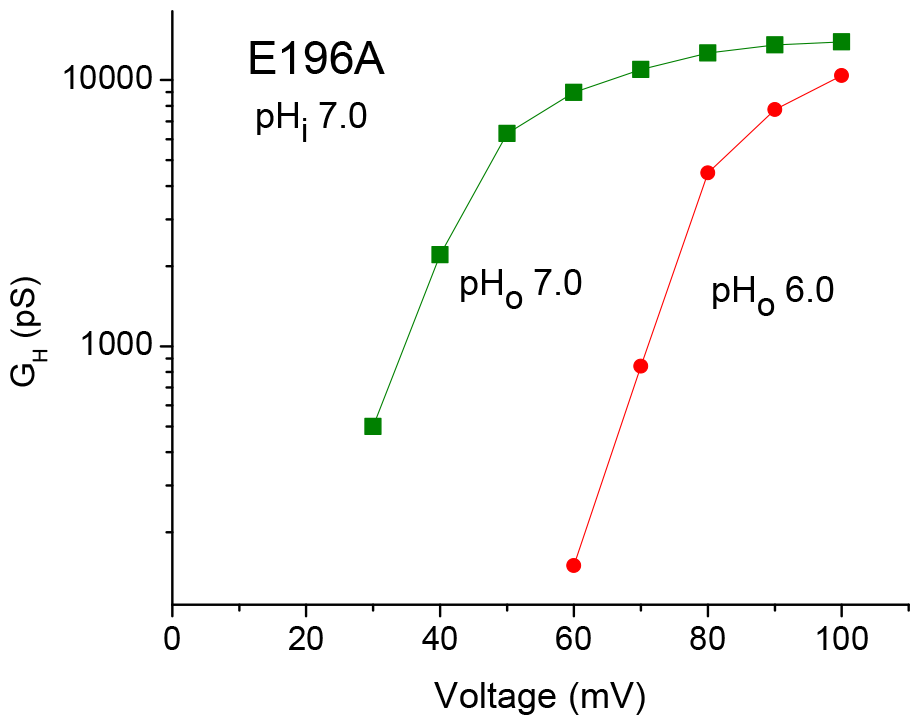
The response of E196A mutant to pH_*o*_ is normal. Proton conductance-voltage relationships at pH_*o*_ 7 (green squares) and pH_*o*_ 6 (red circles) both with pH_*i*_ 7 in a cell transfected with the E196A mutant. The shift of the curves was determined from the voltage at which the G_H_ was 10 % of its maximal value. The mean shift for a change in pH_*o*_ from 7 to 6 was 37.8 ±1.0 mV (mean ±SE) in 3 cells with E196A and 2 with E196Q. For comparison, the shift measured identically in normal hH_v_1 was 38.7 mV (49), and the shift in H168x mutants was 41.4 mV for changes in pH_*o*_ (12) despite the greatly diminished shift for changes in pH_*i*_.

### Three residues may switch protonation state upon depolarization

H_v_1 is gated by changes in voltage or ΔpH across the membrane, and a unique feature of H_v_1 is that the voltage at which the channel opens is dependent on ΔpH (4, 6). Cherny and colleagues formulated a four-state transporter-like model to explain ΔpH-dependent gating, in which one or more titratable residues are alternately exposed to the intracellular or extracellular pH (4, 6). By switching protonation states, these residues can stabilize the closed or open conformation of the channel (4). Thus, residues displaying a large p*K*_a_ shift, especially those changing protonation states between the Hypol and Depol states of H_v_1 may be involved in pH sensing. Although several dozen single mutations failed to attenuate the pH_*o*_-dependence of gating of hH_v_1 (17), H168 mutations to Gln, Ser, or Thr were found to compromise the pH_*i*_ dependence of gating in hH_v_1 (12).

Based on the CpHMD simulations of the Hypol and Depol states of hH_v_1, the p*K*_a_ values of all titratable residues were calculated (Supplemental Table 1). Only three residues, H140, H168, and E196, show a larger than 0.5 pH unit p*K*_a_ difference between the two states that may result in a protonation state change (Figure 2a and b), suggesting that these residues may be involved in sensing pH. Note that 0.5 pH unit reflects the error of the hybrid-solvent CpHMD method (25).

H140 and E196 are both located on the extracellular side, with H140 on S2 and E196 on the loop between S4 and S3 (Figure 2a). However, our analysis showed that neither H140 or E196 is involved in any electrostatic or hydrogen bonding interaction. Instead, the p*K*_a_ change is correlated with the solvent accessibility change. In the Depol state, both H140 and E196 have increased access to water, which results in a p*K*_a_ upshift for H140 and p*K*_a_ downshift for E196. H140 was reported previously not to alter pH_*o*_ sensitivity, with a 46 mV/unit shift for the double mutant H140A/H193A (2, 17). To test if E196 is a pH_*o*_ sensor, we measured the proton conductance-voltage relationships for the E196 mutants. The threshold voltage shifted by an average of 37.8 mV when pH_*o*_ was changed from 6.0 to 7.0 with E196A and E196Q hH_v_1 (Figure 3). As comparison, the shift for the WT hH_v_1 was 38.7 mV (49). Thus, E196 is ruled out as a pH sensor. It is also noteworthy that the Hypol-state p*K*_a_ of E196 has a very large uncertainty. The calculated p*K*_a_’s from the two sets of simulations differ by 2.1 pH units, which is abnormal considering that the typical p*K*_a_ uncertainty from the hybrid-solvent CpHMD simulations is below 0.5 unit (see Supplemental Table 1 and Ref. (25)). This may be attributed to the fact that E196 is at the boundary between the lipid bilayer and solution phase which cannot be accurately treated by the membrane GB model implemented in the membrane-enabled hybrid-solvent CpHMD method (26).

### H168 switches protonation state upon depolarization

H168 which is located on the cytoplasmic side shows a roughly 2 pH-unit p*K*_a_ upshift from 5.5 (5.7/5.4) in the Hypol state to 7.5 (7.7/7.1) in the Depol state, indicating that it changes from the neutral to the charged state upon hH_v_1 activation. The numbers in the parenthesis are p*K*_a_’s from the two set of simulations. We asked why H168 switches its protonation state upon channel activation. H168 is located on the loop at the intracellular end of S3 that connects with S2 (Figure 2a). In the Hypol state, it is largely shielded from solvent as seen from the low hydration number at all pH conditions (Figure 2c); however, in the Depol state, it becomes more exposed to solvent and forms an ion-pair interaction with E171 on the same S3 helix in a pH-dependent manner (Figure 2c). As the simulation pH increases, H168 becomes increasingly deprotonated, and at the same time, the salt bridge fraction of H168–E171 decreases (Figure 2e). Interestingly, the formation of H168–E171 salt bridge is anti-correlated with the salt-bridge interaction between R162 at the intracellular helical end of S2 and E171 (Figure 2e), which becomes increasingly stronger as the simulation pH increases. In other words, R162 competes with H168 for the interaction with E171, and the later is promoted by the protonation of the imidazole group. The probability distribution of the R162–E171 distance offers further support (Supplemental Figure 4). In the Hypol state, the R162–E171 distance has only one peak at ∼3 Å followed by a flat tail at all simulation pH, which indicates that E171 is always in a salt bridge with R162. As a consequence, E171 cannot interact with H168. In the Depol state, however, a minor peak at ∼7.6 Å appears, and its intensity increases as pH decreases while the peak at ∼3 Å lowers with decreasing pH (Supplemental Figure 4). This data is in support of the conclusion that in the Depol state H168 competes with R162 for the interaction with E171, which results in a progressive weakening of the R162–E171 salt-bridge as H168 becomes increasingly protonated or charged.

### S3 helix tilting is correlated with the H168 protonation state change

To further understand the origin of the H168 protonation state change upon hH_v_1 depolarization, we examined possible correlations between the various Hypol-to-Depol conformational changes and the protonation state change of H168. Alignment of the initial Hypol and Depol models shows that the S3 helix is tilted away from the z-axis by about 23° in the Depol relative to the Hypol state (Figure 4a). We wondered if the S3 tilting is correlated with the protonation state of H168. To examine the correlation, we calculated the S3 tilt angle with respect to the principle z-axis for the Hypol and Depol trajectory frames and separated out the data with the protonated or deprotonated H168. If H168 protonation were indeed correlated with S3 tilting, the angle for the frames with the protonated H168 would be larger in both Hypol and Depol state simulations. Figure 4b shows that indeed the tilt angle of S3 is increased from an average of 14.6° with the deprotonated or neutral H168 to an average of 15.5° with the protonated or charged H168. Since H168 is deprotonated in the Hypol state at Ph 7.5 according to the CpHMD data (calculated p*K*_a_ of 5.6, Figure 1b), this data suggests that H168 protonation promotes S3 tilting, which is a conformational hallmark of hH_v_1 activation according to the Depol model of the Tobias and coworkers (23) and consistent with the X-ray crystal structures of the CiVSD channel (PDB 4g7v for the open state and 4g80 for the resting state) (15), which show an increased tilt angle of 14.7° after alignment of both structures using OPM2.

**Figure 4.**
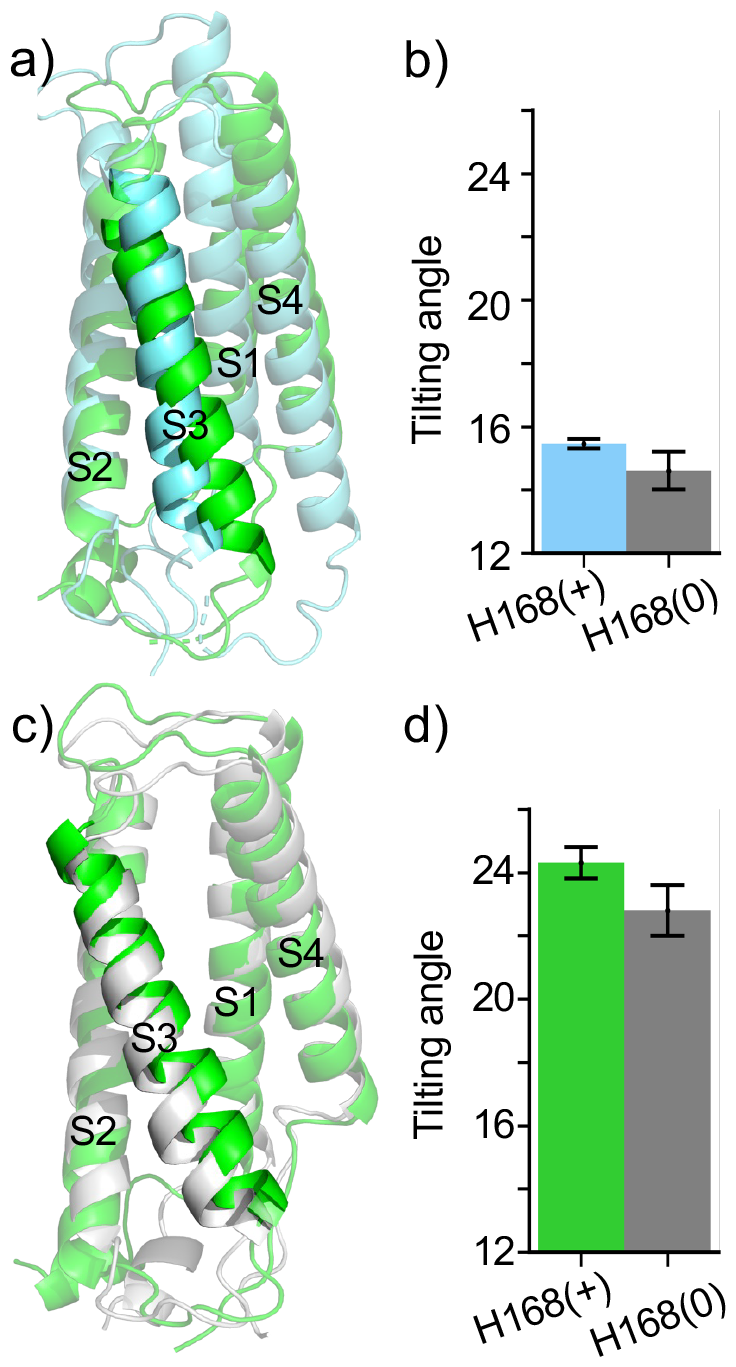
H168 protonation is correlated with the S3 helix tilting. **a)** Alignment of the initial Hypol (blue) and Depol (green) state model obtained from ref(23). **b)** Bar plot of the S3 tilt angle in the Hypol simulations when H168 is protonated (blue) or neutral (grey). A confidence interval of 95% is shown. Replicas at pH 5, 5.5, and 6 (within 0.5 pH unit of the H168 p*K*_a_) were used, with first 10 ns per replica data discarded. The tilt angle is formed by the principle z-axis and the vector defined by backbone atoms of the second first (I172, L173, D174) and last three residues (I186, V187, L188) of the S3 helix. Angle was calculated after structure alignment with the initial structure. **c)** Alignment of the representative snapshots of the Depol state simulation when H168 is protonated (green) or neutral (grey). **d)** Bar plot of the S3 tilt angle in the Depol simulations when H168 is protonated (green) or neutral (grey). Replicas at pH 7, 7.5 and 8 (within 0.5 pH unit of the H168 p*K*_a_) were used, with the first 10 ns per replica data discarded.

Now we consider the Depol state. The simulations showed that the S3 tilt angle is increased from 22.8° when H168 is deprotonated to 24.3° when H168 is protonated (Figure 1c and b). Since H168 spends about 50% of the time sampling the protonated state at pH 7.5 according to the CpHMD data (calculated p*K*_a_ of 7.6 in the Depol state, Figure 1b), this data suggests that H168 deprotonation relaxes the S3 tilting, which is consistent with the conformational change towards the Hypol state. The correlation of H168 protonation with S3 tilting in both Hypol and Depol states suggests that H168 protonation at low pH favors the channel opening, which is consistent with the fact that H_v_1 is activated to extrude protons to the extracellular side at low intracellular pH. In other words, our data and analysis support the hypothesis that H168 acts as a pH_*i*_ sensor (12) and provide mechanistic clues to how this occurs.

## CONCLUDING DISCUSSION

The membrane-enabled hybrid-solvent CpHMD simulations (without an external electric field) were applied to further evaluate the Hypol and Depol models reported by Tobias and coworkers (23) and explore the identities of possible pH sensing residues. The simulations at pH 7.5 showed that the Hypol state is stabilized by two strong and two weak interhelical salt bridges involving the gating charges: the R1–D112 and R2–E153 salt bridges, which are maintained throughout the simulations, and the R1–D185 and R3–D174 salt bridges which are occasionally formed. The only difference from the CiVSD-based closed-state model (16) is the R2–E153 interaction, where R2 is replaced by R3 due to the relatively lower S2 position with respect to S4. The simulations at pH 7.5 showed that the Depol state is stabilized by five salt bridges involving gating charges, between R1 and D123, between R2 and E119/E192, and between R3 and D112/D185. Compared to the Hypol state, D112 changes the salt-bridge partner from R1 to R3, demonstrating that R3 is moved up towards the extracellular side by two clicks in the Depol model (23).

The simulated pH titration based on the Hypol and Depol models of Tobias and coworkers (23) identified three residues with significant p*K*_a_ shifts between the two states that may result in a change in protonation state. These residues are H140, H168, and E196, with calculated p*K*_a_ shifts greater than 1.5 units between the Hypol and Depol states. Among them H140 and E196 are on extracellular side. H140A was shown previously to weaken Zn^2+^ sensitivity (2) but had pH_*o*_ or pH_*i*_ dependent shift in the threshold voltage similar to the WT hH_v_1 (2, 17). E196 is also ruled out as a pH sensor because replacing it with a neutral amino acid (Ala or Gln) did not affect the sensitivity of channel activation to pH_*o*_ (Figure 3).

It is possible that the calculated p*K*_a_ shifts of H140 and E196 are overestimated due to the artifacts of the implicit membrane GB model (27) used in the pH titration. To account for the hydration of the hH_v_1 channel interior (17, 19), a high-dielectric (water) cylinder is implemented in the center of the channel (26). As such, the p*K*_a_ shift accompanying conformational changes for residues near the boundary of the cylinder may be exaggerated. Likewise, the electrostatic energy change for residues moving between the lipid bilayer and solution (such as E196) may also be overestimated.

On the other hand, these results may also indicate that the open-state model we assumed here is not accurate. For example, due to the two-click upward movement of S4, E196 becomes exposed to solvent, which may not be the case if the CiVSD model (16) had been used in the simulations. As discussed above, several types of evidence suggest that the outward displacement of the S4 helix in the open state is less than in the model used here. The apparent agreement of the two-click model with measured gating charge movement was achieved by considering the charge of all of the hH_v_1 monomer basic and acidic side chains (23). This ignores the contribution of obligatory protonation/deprotonation reactions to the measurable gating charge movement (50). Carmona et al. calculated that 60% of the free energy stored in the ΔpH is coupled to voltage sensor activation (35). Another possibility is that our assumption that a pH sensor must exhibit large p*K*_a_ changes during gating may not be correct. Clearly, H_v_1 is strongly modulated by pH_*o*_, and yet the two strongest candidates for sensing pH_*o*_ could be mutated without detectable consequence. It is also conceivable that a pH sensing residue may change its protonation state in the intermediate state (e.g., following the breakage of a salt bridge) and once S4 is up such a change is reverted back (e.g., following the formation of a new salt bridge).

H168 on the intracellular side has been recently suggested as a pH_*i*_ sensor based on the electrophysiology data which showed that mutations of H168 to non-titratable residues Ser, Thr, or Gln significantly weakened the sensitivity of the channel activation to pH_*i*_ (12). The calculated p*K*_a_ value of H168 is 5.6 in the Hypol and 7.6 in the Depol state, indicating that H168 can switch from being deprotonated or neutral to being protonated or charged upon channel depolarization. The analysis showed that the p*K*_a_ shift of H168 can be attributed to a conformational change in going from the Hypol to the Depol state, in which H168 forms a salt bridge with E171 of S3 in a pH-dependent manner. Interestingly, H168 competes with R162 at the intracellular end of S2 for the interaction with E171. The suggestion that H168 is a pH_*i*_ sensor is further supported by the correlation between H168 protonation and increased S3 tilting in the simulations, while S3 tilting is a conformational hallmark of the Depol model of hH_v_1 (23) demonstrated by the X-ray structure of open-state CiVSD (15). Thus, both the protonation state switch of H168 and its correlation with the depolarization induced conformational change of hH_v_1 support the role of H168 as the pH_*i*_ sensor.

We compare the Hypol state p*K*_a_’s of H140 (5.7/5.7), H168 (5.6/5.4), and E196 (7.4/5.3) to the recent estimates by Jardin et al. (22), who used hybrid-solvent Monte-Carlo constant pH MD simulations (51) starting from a homology model (21) based on the resting state crystal structure of CiVSD (15). Their simulations gave the p*K*_a_’s of 5.35, 6.16, and 8.34 for H140, H168, and E196, respectively. Despite the use of the different resting state models and the different implicit-solvent model as well as the neglect of membrane in the simulations of Jardin et al. (22), the calculated p*K*_a_’s of the three residues are in qualitative agreement. That is, the p*K*_a_’s of H140 and H168 are downshifted and the p*K*_a_ of E196 is upshifted relative to the respective model p*K*_a_ values, consistent with our analysis that all three residues are at least partially buried in the Hypol state.

The above agreement is encouraging and suggests that the uncertainty in the computational identification of potential pH sensors through the calculation of p*K*_a_ shifts may lie primarily on the Depol state model used in the simulations. We envisage future studies employing the more accurate structure models and the recently developed GPU-accelerated all-atom particle Ewald CpHMD simulation technique (52, 53) will improve the understanding of pH sensors and unveil the pH-dependent gating mechanism of hH_v_1.

## MATERIALS AND METHODS

### Simulation methods and protocols

The three-stage protocol for CpHMD simulations of transmembrane proteins described in Ref (54) was followed. Below we give system specific information, and more details can be found in Ref (54).

#### System preparation

The simulation starting structures were the hyperpolarized (Hypol) and depolarized (Depol) models of hH_v_1 obtained by Tobias and coworkers using the conventional fixed-protonation-state MD simulations under a constant uniform electric field which corresponds to a transmembrane voltage of -150 mV for the Hypol and 150 mV for the Depol state (23). The Membrane Builder module of CHARMM-GUI interface (55), which employs the CHARMM program (56), was used to assemble the simulation system. The truncated N- (F88) and C- (R230) terminals were acetylated (CH_3_CO) and amidated (NH_2_), respectively. In the system preparation and membrane equilibration stages, the default protonation states Asp(-)/Glu(-)/Lys(+)/Arg(+)/Cys(0)/His(0, HSD) were used. The protein was inserted in a pre-assembled bilayer of 1-palmitoyl-2-oleoyl-sn-glycero-3-phosphocholine (POPC) lipids. The upper and lower leaflets contain 84/83 and 81/80 lipids, respectively, for the Hypol/Depol state. A water layer of 15 Å thickness was added to both sides of the lipid bilayer, and 15/17 water molecules were placed in the channel pore for the Hypol/Depol state. To neutralize the simulation system (with the default protonation states) and reach the physiological ionic strength of 0.15 M, 24 sodium and 33 chloride ions for the Hypol state or 21 sodium and 30 chloride ions for the Depol state were added. The final systems comprised about 50,000 atoms.

#### Membrane equilibration

The CHARMM22/CMAP force field was employed to describe the protein (57, 58), while lipids and ions were described by the CHARMM36 lipid force field (59). The CHARMM modified TIP3P model was used to represent water (58). The positions of lipids were relaxed with the proton heavy atom restrained following the multi-step protocol in CHARMM-GUI (55). After the initial relaxation, the protein-membrane complex was subject to a 200-ns conventional fixed-protonation-state MD simulation using the OpenMM program (60). The simulation was performed in the NPT ensemble where the protein heavy atoms were restrained with a harmonic force constant of 1 kcal/mol/Å^2^. The Hoover thermostat (61) and the Langevin piston coupling method (62) were employed to maintain the temperature of 310 K and the pressure of 1 atm, respectively. The particle mesh Ewald method was used to calculate long-range electrostatics (63) with a real-space cutoff of 12 Å and the reciprocal space calculation used a 1-Å grid spacing and sixth-order spline interpolation. The SHAKE algorithm was used to constrain bonds involving hydrogen to enable a time step of 2 fs. The area per lipid (APL) and lipid order parameter were used to assess convergence of the lipid bilayer equilibration. More details and MD settings are given in Ref (54).

#### CpHMD simulations and analysis

Following the membrane equilibration, dummy hydrogen atoms were added to the carboxylate groups of Asp and Glu and the singly protonated histidine HSD is replaced with HSP (the doubly protonated histidine). The membrane-enabled hybrid-solvent CpHMD simulations were performed using the CHARMM package (version c45a1) (56). In the membrane GBSW model (27), the low-dielectric region had a thickness of 33 Å and the radius of the high-dielectric cylinder that encompasses the channel was 12 Å. The MMFP facility in CHARMM was employed to restrain the center of mass of the protein with a harmonic force constant of 1.0 kcal/mol/Å^2^. During the CpHMD simulation, all Asp, Glu, and His sidechains were allowed to titrate, while Lys and Arg sidechains were fixed in the protonated or charged state. The single-pH (pH=7) CpHMD simulation was first conducted to relax the protein heavy atoms by gradually releasing the restraint potential within 1 ns (detailed protocol given in Ref (54)). Next, the restraints were removed and the protein-membrane system was further equilibrated for 1 ns. In the production run, the pH replica-exchange protocol was employed to accelerate the sampling convergence of the coupled protonation and conformational dynamics (25). In Run 1, the protocol used a total of 16 pH replicas in the pH range of 3 to 11.5 with an interval of 0.5 pH unit. The simulation for each replica lasted 25 ns, with the aggregate sampling time of 400 ns. In Run 2, the protocol used 24 replicas in the pH range 1 to 8 with an interval of 0.25 pH unit except for the pH ranges 1–2.5 and 7–8 where an interval of 0.5 pH unit was used. Each replica was simulated for 30 ns and the aggregate sampling time was 720 ns. Adjacent replicas were allowed to exchange every 500 MD steps or 1 ps.

The p*K*_a_ value of a specific residue was calculated by fitting the unprotonated fraction (*S*) to the generalized Henderson-Hasselbalch equation,

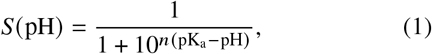

where *n* is the Hill coefficient.

The software Cpptraj (64) was used for trajectory analysis (salt bridge distance and helix tilting angle calculation, etc.). Trajectory visualization was performed using VMD (65) and snapshot rendering was produced using Pymol (66). All analysis was based on the trajectories after discarding the first 10 ns per pH replica.

### Experimental methods

The materials and procedures used for generating mutants, transfection, tight-seal voltage clamp recording, and data analysis are as described previously (18).

## STATEMENT OF COMPETING INTERESTS

The authors declare no competing interests.

## ACKNOWLEDGMENTS

We thank I. Scott Ramsey for the helpful discussion. J.S. and M. S. were supported by the National Institutes of Health (R35GM148261). Funding for Y.H. and Z.C. is provided by the National Natural Science Foundation of China (11804114) and the Natural Science Foundation of Fujian Province (2023J01329). T.E.D. and V.V.C. were supported by the National Institutes of Health (R35GM151963).

## SUPPLEMENTARY MATERIAL

An online supplement to this article can be found by visiting BJ Online at http://www.biophysj.org.

## Notes

### Competing Interest Statement

The authors have declared no competing interest.

### Summary of Updates

Electrophysiological experiment data were added and the present manuscript focuses on the discussion of H168 identified as the interior pH sensing residue.

## REFERENCES

1. Thomas, R. C., and R. W. Meech, 1982. Hydrogen Ion Currents and Intracellular pH in Depolarized Voltage-Clamped Snail Neurones. Nature 299:826–828.

2. Ramsey, I. S., M. M. Moran, J. A. Chong, and D. E. Clapham, 2006. A Voltage-Gated Proton-Selective Channel Lacking the Pore Domain. Nature 440:1213.

3. Sasaki, M., M. Takagi, and Y. Okamura, 2006. A Voltage Sensor-Domain Protein Is a Voltage-Gated Proton Channel. Science 312:589–592.

4. Cherny, V. V., V. S. Markin, and T. E. DeCoursey, 1995. The Voltage-Activated Hydrogen Ion Conductance in Rat Alveolar Epithelial Cells Is Determined by the pH Gradient. J. Gen. Physiol. 105:861–896.

5. Musset, B., S. M. E. Smith, S. Rajan, D. Morgan, V. V. Cherny, and T. E. DeCoursey, 2011. Aspartate 112 Is the Selectivity Filter of the Human Voltage-Gated Proton Channel. Nature 480:273–277.

6. DeCoursey, T. E., 2018. Voltage and pH Sensing by the Voltage-Gated Proton Channel, HV1. J. R. Soc. Interface 15.

7. He, J., R. M. Ritzel, and J. Wu, 2021. Functions and Mechanisms of the Voltage-Gated Proton Channel Hv1 in Brain and Spinal Cord Injury. Front. Cell. Neurosci. 15:662971.

8. Tombola, F., M. H. Ulbrich, and E. Y. Isacoff, 2008. The Voltage-Gated Proton Channel Hv1 Has Two Pores, Each Controlled by One Voltage Sensor. Neuron 58:546–556.

9. Lee, S.-Y., J. A. Letts, and R. MacKinnon, 2008. Dimeric Subunit Stoichiometry of the Human Voltage-Dependent Proton Channel Hv1. Proc. Natl. Acad. Sci. U.S.A. 105:7692–7695.

10. Koch, H. P., T. Kurokawa, Y. Okochi, M. Sasaki, Y. Okamura, and H. P. Larsson, 2008. Multimeric Nature of Voltage-Gated Proton Channels. Proc. Natl. Acad. Sci. U.S.A. 105:9111–9116.

11. DeCoursey, T. E., D. Morgan, B. Musset, and V. V. Cherny, 2016. Insights into the Structure and Function of HV1 from a Meta-Analysis of Mutation Studies. J. Gen. Physiol. 148:97–118.

12. Cherny, V. V., D. Morgan, S. Thomas, S. M. Smith, and T. E. DeCoursey, 2018. Histidine-168 Is Crucial for ΔpH-dependent Gating of the Human Voltage-Gated Proton Channel, hHv1. J. Gen. Physiol. 150:851–862.

13. Takeshita, K., S. Sakata, E. Yamashita, Y. Fujiwara, Kawanabe, T. Kurokawa, Y. Okochi, M. Matsuda, H. Narita, Y. Okamura, and A. Nakagawa, 2014. X-Ray Crystal Structure of Voltage-Gated Proton Channel. Nat. Struct. Mol. Biol. 21:352–357.

14. Bayrhuber, M., I. Maslennikov, W. Kwiatkowski, Sobol, C. Wierschem, C. Eichmann, L. Frey, and R. Riek, 2019. Nuclear Magnetic Resonance Solution Structure and Functional Behavior of the Human Proton Channel. Biochemistry 58:4017–4027.

15. Li, Q., S. Wanderling, M. Paduch, D. Medovoy, A. Singharoy, R. McGreevy, C. A. Villalba-Galea, R. E. Hulse, B. Roux, K. Schulten, A. Kossiakoff, and E. Perozo, 2014. Structural Mechanism of Voltage-Dependent Gating in an Isolated Voltage-Sensing Domain. Nat. Struct. Mol. Biol. 21:244–252.

16. Li, Q., R. Shen, J. S. Treger, S. S. Wanderling, W. Milewski, K. Siwowska, F. Bezanilla, and E. Perozo, 2015. Resting State of the Human Proton Channel Dimer in a Lipid Bilayer. Proc. Natl. Acad. Sci. U.S.A. 112:E5926–E5935.

17. Ramsey, I. S., Y. Mokrab, I. Carvacho, Z. A. Sands, M. S. P. Sansom, and D. E. Clapham, 2010. An Aqueous H+ Permeation Pathway in the Voltage-Gated Proton Channel Hv1. Nat. Struct. Mol. Biol. 17:869–875.

18. Kulleperuma, K., S. M. Smith, D. Morgan, B. Musset, J. Holyoake, N. Chakrabarti, V. V. Cherny, T. E. DeCoursey, and R. Pomès, 2013. Construction and Validation of a Homology Model of the Human Voltage-Gated Proton Channel hHV1. J. Gen. Physiol. 141:445–465.

19. Gianti, E., L. Delemotte, M. L. Klein, and V. Carnevale, 2016. On the Role of Water Density Fluctuations in the Inhibition of a Proton Channel. Proc. Natl. Acad. Sci. U.S.A. 113:E8359–E8368.

20. Randolph, A. L., Y. Mokrab, A. L. Bennett, M. S. Sansom, and I. S. Ramsey, 2016. Proton Currents Constrain Structural Models of Voltage Sensor Activation. eLife 5:e18017.

21. Jardin, C., G. Chaves, and B. Musset, 2020. Assessing Structural Determinants of Zn2+ Binding to Human HV1 via Multiple MD Simulations. Biophys. J. 118:1221– 1233.

22. Jardin, C., N. Ohlwein, A. Franzen, G. Chaves, and B. Musset, 2022. The pH-dependent Gating of the Human Voltage-Gated Proton Channel from Computational Simulations. Phys. Chem. Chem. Phys. 24:9964–9977.

23. Geragotelis, A. D., M. L. Wood, H. Göddeke, L. Hong, P. D. Webster, E. K. Wong, J. A. Freites, F. Tombola, and D. J. Tobias, 2020. Voltage-Dependent Structural Models of the Human Hv1 Proton Channel from Long-Timescale Molecular Dynamics Simulations. Proc. Natl. Acad. Sci. U.S.A. 117:13490–13498.

24. Banh, R., V. V. Cherny, D. Morgan, B. Musset, S. Thomas, K. Kulleperuma, S. M. E. Smith, R. Pomès, and T. E. DeCoursey, 2019. Hydrophobic Gasket Mutation Produces Gating Pore Currents in Closed Human Voltage-Gated Proton Channels. Proc. Natl. Acad. Sci. U.S.A. 116:18951– 18961.

25. Wallace, J. A., and J. K. Shen, 2011. Continuous Constant pH Molecular Dynamics in Explicit Solvent with pH-Based Replica Exchange. J. Chem. Theory Comput. 7:2617–2629.

26. Huang, Y., W. Chen, D. L. Dotson, O. Beckstein, and J. Shen, 2016. Mechanism of pH-dependent Activation of the Sodium-Proton Antiporter NhaA. Nat. Commun. 7:12940.

27. Im, W., M. Feig, and C. L. Brooks, III, 2003. An Implicit Membrane Generalized Born Theory for the Study of Structure, Stability, and Interactions of Membrane Proteins. Biophys. J. 85:2900–2918.

28. Yue, Z., W. Chen, H. I. Zgurskaya, and J. Shen, 2017. Constant pH Molecular Dynamics Reveals How Proton Release Drives the Conformational Transition of a Transmembrane Efflux Pump. J. Chem. Theory Comput. 13:6405–6414.

29. Henderson, J. A., Y. Huang, O. Beckstein, and J. Shen, 2020. Alternative Proton-Binding Site and Long-Distance Coupling in Escherichia Coli Sodium–Proton Antiporter NhaA. Proc. Natl. Acad. Sci. U.S.A. 117:25517–25522.

30. Chen, W., Y. Huang, and J. Shen, 2016. Conformational Activation of a Transmembrane Proton Channel from Constant pH Molecular Dynamics. J. Phys. Chem. Lett. 7:3961–3966.

31. Yarov-Yarovoy, V., P. G. DeCaen, R. E. Westenbroek, C.-Y. Pan, T. Scheuer, D. Baker, and W. A. Catterall, 2012. Structural Basis for Gating Charge Movement in the Voltage Sensor of a Sodium Channel. Proceedings of the National Academy of Sciences 109.

32. Morgan, D., B. Musset, K. Kulleperuma, S. M. Smith, S. Rajan, V. V. Cherny, R. Pomès, and T. E. DeCoursey, 2013. Peregrination of the Selectivity Filter Delineates the Pore of the Human Voltage-Gated Proton Channel hH V 1. J. Gen. Physiol. 142:625–640.

33. Gonzalez, C., H. P. Koch, B. M. Drum, and H. P. Larsson, 2010. Strong Cooperativity between Subunits in Voltage-Gated Proton Channels. Nat. Struct. Mol. Biol. 17:51–56.

34. Gonzalez, C., S. Rebolledo, M. E. Perez, and H. P. Larsson, 2013. Molecular Mechanism of Voltage Sensing in Voltage-Gated Proton Channels. J. Gen. Physiol. 141:275– 285.

35. Carmona, E. M., M. Fernandez, J. J. Alvear-Arias, Neely, H. P. Larsson, O. Alvarez, J. A. Garate, R. Latorre, and C. Gonzalez, 2021. The Voltage Sensor Is Responsible for ΔpH Dependence in H v 1 Channels. Proc. Natl. Acad. Sci. U.S.A. 118:e2025556118.

36. Huxley, A. F., and R. Stämpfli, 1951. Direct Determination of Membrane Resting Potential and Action Potential in Single Myelinated Nerve Fibres. J. Physiol. 112:476– 495.

37. Lewis, R., K. E. Asplin, G. Bruce, C. Dart, A. Mobasheri, and R. Barrett-Jolley, 2011. The Role of the Membrane Potential in Chondrocyte Volume Regulation. J. Cell. Physiol. 226:2979–2986.

38. DeCoursey, T. E., 2000. Hypothesis: Do Voltage-Gated H + Channels in Alveolar Epithelial Cells Contribute to CO2 Elimination by the Lung? Am. J. Physiol. 278:C1–C10.

39. Jankowski, A., and S. Grinstein, 1999. A Noninvasive Fluorimetric Procedure for Measurement of Membrane Potential. J. Biol. Chem. 274:26098–26104.

40. Murphy, R., and T. E. DeCoursey, 2006. Charge Compensation during the Phagocyte Respiratory Burst. Biochim. Biophys. Acta 1757:996–1011.

41. Chernomordik, L., S. Sukharev, S. Popov, V. Pastushenko, Sokirko, I. Abidor, and Y. Chizmadzhev, 1987. The Electrical Breakdown of Cell and Lipid Membranes: The Similarity of Phenomenologies. Biochim. Biophys. Acta 902:360–373.

42. DeCoursey, T., and V. Cherny, 1994. Voltage-Activated Hydrogen Ion Currents. J. Membr. Biol. 141.

43. Villalba-Galea, C. A., 2014. Hv1 Proton Channel Opening Is Preceded by a Voltage-independent Transition. Biophys. J. 107:1564–1572.

44. Carmona, E. M., H. P. Larsson, A. Neely, O. Alvarez, R. Latorre, and C. Gonzalez, 2018. Gating Charge Displacement in a Monomeric Voltage-Gated Proton (H v 1) Channel. Proceedings of the National Academy of Sciences 115:9240–9245.

45. Cherny, V. V., B. Musset, D. Morgan, S. Thomas, S. M. Smith, and T. E. DeCoursey, 2020. Engineered High-Affinity Zinc Binding Site Reveals Gating Configurations of a Human Proton Channel. J. Gen. Physiol. 152:e202012664.

46. Decoursey, T. E., 2003. Voltage-Gated Proton Channels and Other Proton Transfer Pathways. Physiol. Rev. 83:475– 579.

47. Musset, B., S. M. E. Smith, S. Rajan, V. V. Cherny, S. Sujai, D. Morgan, and T. E. DeCoursey, 2010. Zinc Inhibition of Monomeric and Dimeric Proton Channels Suggests Cooperative Gating. J. Physiol. 588:1435–1449.

48. Fujiwara, Y., T. Kurokawa, K. Takeshita, M. Kobayashi, Y. Okochi, A. Nakagawa, and Y. Okamura, 2012. The Cytoplasmic Coiled-Coil Mediates Cooperative Gating Temperature Sensitivity in the Voltage-Gated H+ Channel Hv1. Nat. Commun. 3:816.

49. Cherny, V. V., D. Morgan, B. Musset, G. Chaves, S. M. Smith, and T. E. DeCoursey, 2015. Tryptophan 207 Is Crucial to the Unique Properties of the Human Voltage-Gated Proton Channel, hHV1. J. Gen. Physiol. 146:343– 356.

50. Sokolov, V. S., V. V. Cherny, A. G. Ayuyan, and T. E. DeCoursey, 2021. Analysis of an Electrostatic Mechanism for ΔpH Dependent Gating of the Voltage-Gated Proton Channel, HV1, Supports a Contribution of Protons to Gating Charge. Biochim. Biophys. Acta 1862:148480.

51. Swails, J. M., D. M. York, and A. E. Roitberg, 2014. Constant pH Replica Exchange Molecular Dynamics in Explicit Solvent Using Discrete Protonation States: Implementation, Testing, and Validation. J. Chem. Theory Comput. 10:1341–1352.

52. Huang, Y., W. Chen, J. A. Wallace, and J. Shen, 2016. All-Atom Continuous Constant pH Molecular Dynamics With Particle Mesh Ewald and Titratable Water. J. Chem. Theory Comput. 12:5411–5421.

53. Harris, J. A., R. Liu, V. Martins de Oliveira, E. A. Vázquez-Montelongo, J. A. Henderson, and J. Shen, 2022. GPU-Accelerated All-Atom Particle-Mesh Ewald Continuous Constant pH Molecular Dynamics in Amber. J. Chem. Theory Comput. 18:7510–7527.

54. Huang, Y., J. A. Henderson, and J. Shen, 2021. Continuous Constant pH Molecular Dynamics Simulations of Transmembrane Proteins. In Methods in Molecular Biology, Springer, New York, volume 2302 of Structure and Function of Membrane Proteins, 275–287.

55. Jo, S., T. Kim, V. G. Iyer, and W. Im, 2008. CHARMM-GUI: A Web-Based Graphical User Interface for CHARMM. J. Comput. Chem. 29:1859–1865.

56. Brooks, B., C. Brooks, A. MacKerell, L. Nilsson, R. Petrella, B. Roux, Y. Won, G. Archontis, C. Bartels, S. Boresch, A. Caflisch, L. Caves, Q. Cui, A. Dinner, M. Feig, S. Fischer, J. Gao, M. Hodoscek, W. Im, K. Kuczera, T. Lazaridis, J. Ma, V. Ovchinnikov, E. Paci, R. Pastor, C. Post, J. Pu, M. Schaefer, B. Tidor, R. M. Venable, H. L. Woodcock, X. Wu, W. Yang, D. York, and M. Karplus, 2009. CHARMM: The Biomolecular Simulation Program. J. Comput. Chem. 30:1545–1614.

57. MacKerell, A. D., D. Bashford, M. Bellott, R. L. Dunbrack, J. D. Evanseck, M. J. Field, S. Fischer, J. Gao, H. Guo, S. Ha, D. Joseph-McCarthy, L. Kuchnir, K. Kuczera, F. T. K. Lau, C. Mattos, S. Michnick, T. Ngo, D. T. Nguyen, B. Prodhom, W. E. Reiher, B. Roux, M. Schlenkrich, J. C. Smith, R. Stote, J. Straub, M. Watanabe, J. Wiórkiewicz-Kuczera, D. Yin, and M. Karplus, 1998. All-Atom Empirical Potential for Molecular Modeling and Dynamics Studies of Proteins. J. Phys. Chem. B 102:3586–3616.

58. Mackerell, A. D., M. Feig, and C. L. Brooks, 2004. Extending the Treatment of Backbone Energetics in Protein Force Fields: Limitations of Gas-Phase Quantum Mechanics in Reproducing Protein Conformational Distributions in Molecular Dynamics Simulations. J. Comput. Chem. 25:1400–1415.

59. Klauda, J. B., R. M. Venable, J. A. Freites, J. W. O’Connor, D. J. Tobias, C. Mondragon-Ramirez, I. Vorobyov, A. D. MacKerell, and R. W. Pastor, 2010. Update of the CHARMM All-Atom Additive Force Field for Lipids: Validation on Six Lipid Types. J. Phys. Chem. B 114:7830– 7843.

60. Eastman, P., J. Swails, J. D. Chodera, R. T. McGibbon, Y. Zhao, K. A. Beauchamp, L.-P. Wang, A. C. Simmonett, M. P. Harrigan, C. D. Stern, R. P. Wiewiora, B. R. Brooks, and V. S. Pande, 2017. OpenMM 7: Rapid Development of High Performance Algorithms for Molecular Dynamics. PLoS Comput. Biol. 13:e1005659.

61. Hoover, W. G., 1985. Canonical Dynamics: Equilibration Phase-Space Distributions. Phys. Rev. A 31:1695–1697.

62. Feller, S. E., Y. Zhang, R. W. Pastor, and B. R. Brooks, 1995. Constant Pressure Molecular Dynamics Simulation: The Langevin Piston Method. J. Chem. Phys. 103:4613– 4621.

63. Darden, T., D. York, and L. Pedersen, 1993. Particle Mesh Ewald: An W Log(N) Method for Ewald Sums in Large Systems. J. Chem. Phys. 98:4.

64. Roe, D. R., and T. E. Cheatham, 2013. PTRAJ and CPPTRAJ: Software for Processing and Analysis of Molecular Dynamics Trajectory Data. Journal of Chemical Theory and Computation 9:3084–3095.

65. Humphrey, W., A. Dalke, and K. Schulten, 1996. VMD: Visual Molecular Dynamics. Journal of Molecular Graphics 14:33–38.

66. Schrödinger, LLC, 2015. The PyMOL Molecular Graphics System, Version 1.8.

