## Supporting Information for "Interior pH Sensing Residue of Human Voltage-Gated Proton Channel H_v_1 is Histidine 168"

Manuscript submitted to **Biophysical** *Journal*

Article

### SUPPLEMENTAL TABLE

Table 1: The calculated p$K_a$'s titratable residues in the Hypol and Depol states of hH$_v$1 and the p$K_a$ shifts between the two states

| ASP | | | | GLU | | | |
|---|---|---|---|---|---|---|---|
| Residue | Hypol | Depol | Δp$K_a$ | Residue | Hypol | Depol | Δp$K_a$ |
| D112 | 3.5 (3.6) | 2.2 (3.2) | 1.3 (0.4) | E119 | 4.3 (4.4) | 4.8 (4.7) | -0.5 (-0.3) |
| D123 | 3.7 (3.7) | 3.3 (3.1) | 0.4 (0.6) | E153 | 3.6 (3.4) | 2.9 (3.7) | 0.7 (-0.3) |
| D130 | 3.4 (3.5) | 3.5 (3.3) | -0.1 (0.2) | E164 | 3.1 (3.4) | 2.5 (2.8) | 0.6 (0.6) |
| D174 | 0.1 (1.1) | 1.5 (2.0) | -1.4 (-0.9)* | E171 | 2.9 (3.1) | 2.8 (2.9) | 0.1 (0.2) |
| D185 | 3.8 (4.2) | 3.3 (3.6) | 0.5 (0.6) | E192 | 3.8 (3.7) | 2.5 (3.2) | 1.3 (0.5) |
| | | | | **E196** | 7.4 (5.3) | 4.1 (4.0) | 3.3 (1.3) |
| | | | | E225 | 3.5 (3.8) | 3.5 (3.5) | 0.0 (0.3) |
| HIS | | | | | | | |
| Residue | Hypol | Depol | Δp$K_a$ | | | | |
| H99 | 6.0 (6.2) | 6.0 (6.4) | 0.0 (-0.2) | | | | |
| **H140** | 5.7 (5.7) | 7.2 (6.9) | -1.5 (-1.2) | | | | |
| H167 | 7.5 (7.3) | 7.4 (7.3) | 0.1 (0.0) | | | | |
| **H168** | 5.6 (5.4) | 7.6 (7.1) | -2.0 (-1.7) | | | | |
| H193 | 8.4 (8.4) | 8.6 (8.6) | -0.2 (-0.2) | | | | |

The p$K_a$'s and p$K_a$ shifts calculated from the second set 2 are listed in parentheses. Residues in bold show p$K_a$ shifts may result in a protonation state change at pH 7 as discussed in the main text. * The p$K_a$'s and p$K_a$ shifts are highly approximate due to the incomplete titration.

### SUPPLEMENTAL FIGURES

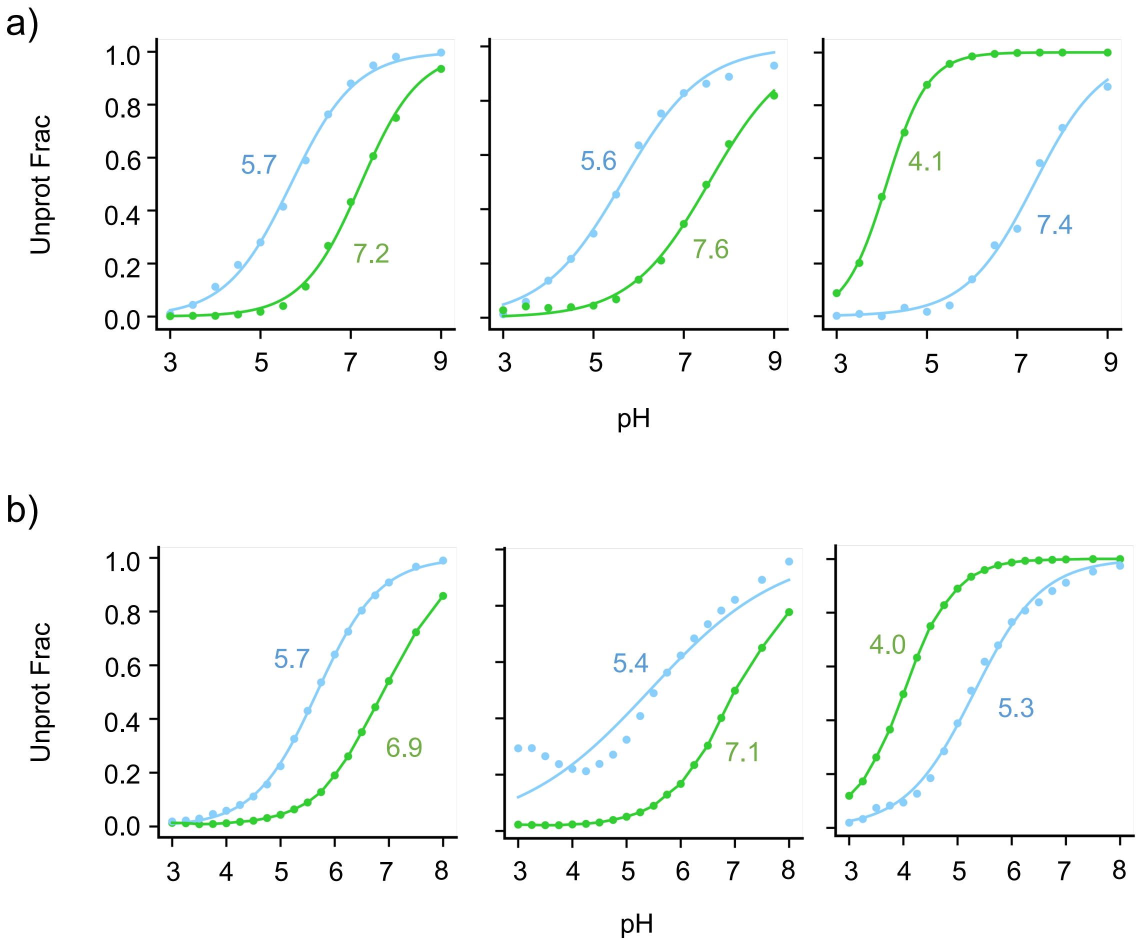


Figure 1: **Titration plots for H140, H168, and E196.** The deprotonated fraction is calculated at different simulation pH for simulation set 1 (a) and set 2 (b). The data of the Hypol and Depol states are shown in blue and green, respectively. The best fits to the generalized Henderson-Hasselbalch equation are shown in solid curves. The estimated p$K_a$'s are given. Data from the pH conditions below 3 and above 9 are hidden.

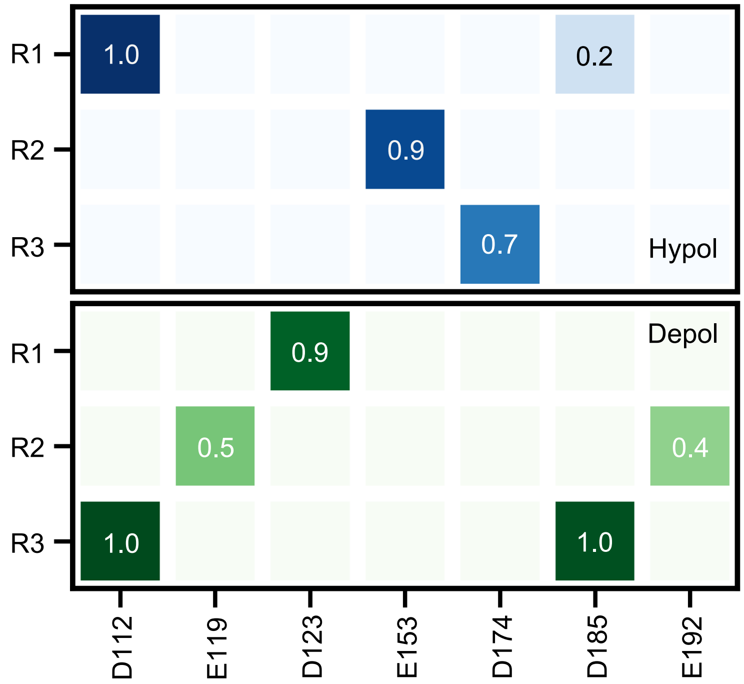


Figure 2: **The interhelical salt bridges in the Hypol and Depol states of hH$_v$1 calculated from the second set of replica-exchange CpHMD simulations.** Comparison of the salt-bridge occupancies (probabilities) in the Hypol (top) and Depol (bottom) states from the second set of simulations of hH$_v$1 at pH 7.5. A salt bridge is considered formed when the minimum distance between the carboxylate oxygen and the arginine guanidinium nitrogen is below 3.5 Å.

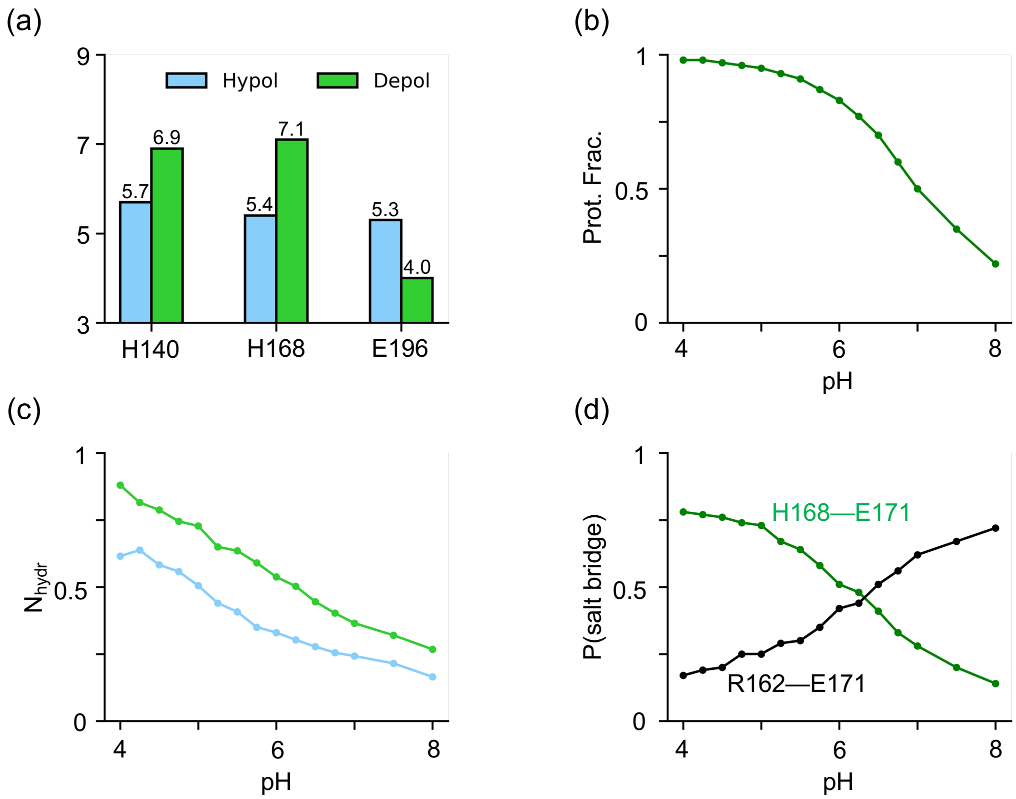


Figure 3: **Why does H168 switch protonation state? Results from the second set of replica-exchange CpHMD simulations.** **a)** The calculated p$K_a$ values of H140, H168, and E196 in the Hypol (blue) and Depol (green) states of $hH_v1$. These are the only residues that show a larger than 0.5-unit p$K_a$ difference between the two states. **b)** Protonation fraction of H168 in the Depol state at different simulation pH. **c)** Number of water (hydration number) near H168, calculated as the number of water molecules within 3.4 Å from H168. **d)** Probabilities (or occupancies) of the H168–E171 (green) and R162–E171 (black) salt bridges in the Depol state at different simulation pH.

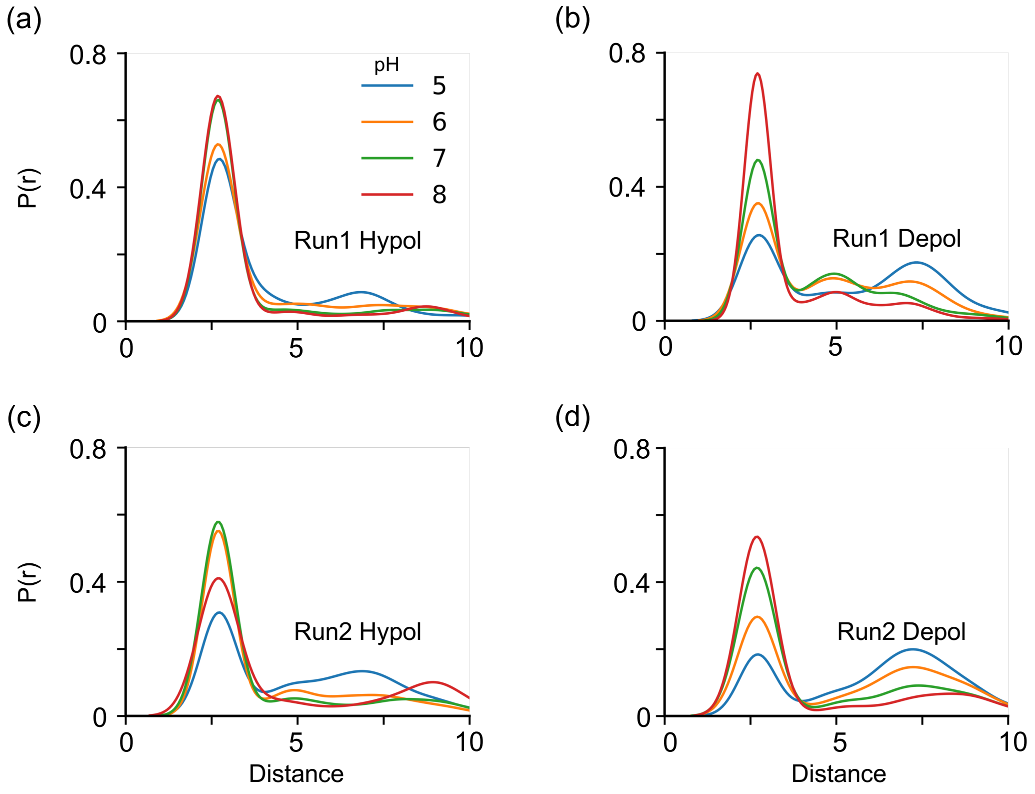


Figure 4: **The pH-dependent interaction between R162 and E171 from both sets of replica-exchange CpHMD simulation in the Hypol and Depol states.** The minimum distance between the Arg side chain nitrogen and the Glu side chain oxygens are plotted for both sets of simulations at the simulation pH at 5, 6, 7, and 8 after discarding the first 10 ns/replica data.

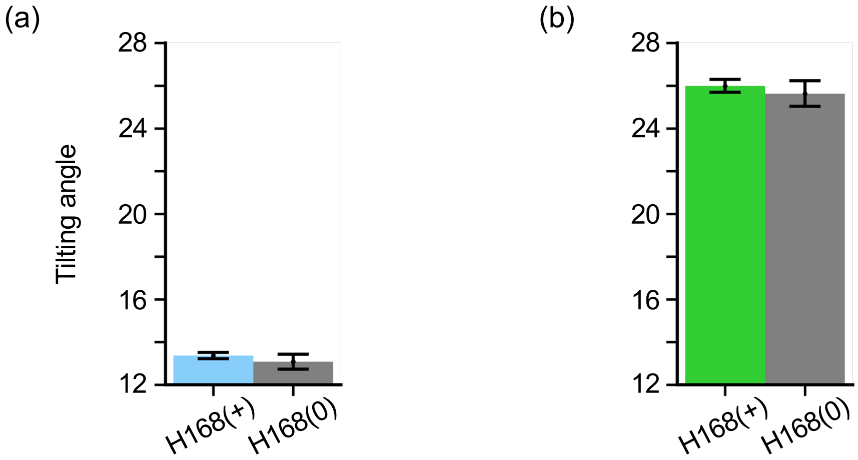


Figure 5: **H168 protonation is correlated with the S3 helix tilting (Run2). a)** Bar plot of the S3 tilt angle in the Hypol simulations when H168 is protonated (blue) or neutral (grey) in Run2. A confidence interval of 95% is shown. Replicas at pH 5, 5.5, and 6 (within 0.5 pH unit of the H168 p$K_a$ (5.4 in Run2)) were used, with the first 10 ns per replica data discarded. The tilt angle is formed by the principle z-axis and the vector defined by backbone atoms of the second first (I172, L173, D174) and last three residues (I186, V187, L188) of the S3 helix. Angle was calculated after structure alignment with the initial structure. **b)** Bar plot of the S3 tilt angle in the Depol simulations when H168 is protonated (green) or neutral (grey) in Run2. Replicas at pH 6.5, 7, and 7.5 (within 0.5 pH unit of the H168 p$K_a$ (7.1 in Run2)) were used, with the first 10 ns per replica data discarded.
